# Identification of endogenous Adenomatous polyposis coli interaction partners and β-catenin-independent targets by proteomics

**DOI:** 10.1101/258400

**Authors:** Olesja Popow, Michael H. Tatham, João A. Paulo, Alejandro Rojas-Fernandez, Nicolas Loyer, Ian P. Newton, Jens Januschke, Kevin M. Haigis, Inke Näthke

## Abstract

*Adenomatous polyposis coli* (*APC*) is the most frequently mutated gene in colorectal cancer. APC negatively regulates the pro-proliferative Wnt signaling pathway by promoting the degradation of β-catenin, but the extent to which APC exerts Wnt/β-catenin-independent tumor suppressive activity is unclear. To identify interaction partners and β-catenin-independent targets of endogenous, full-length APC, we applied label-free and multiplexed TMT mass spectrometry. Affinity enrichment-mass spectrometry revealed over 150 previously unidentified APC interaction partners. Moreover, our global proteomic analysis revealed that roughly half of the protein expression changes that occur in response to APC loss are independent of β-catenin. By combining these two analyses, we identified Misshapen-like kinase 1 (MINK1) as a putative substrate of an alternative APC-containing destruction complex and provide evidence for the potential contribution of MINK1 to *APC* mutant phenotypes. Collectively, our results highlight the extent and importance of Wnt-independent APC functions in epithelial biology and disease.

## Introduction

Mutations in *Adenomatous polyposis coli* (*APC*) are a frequent (> 80%) and early event in the development of sporadic colorectal cancer (CRC; Miyoshi et al., 1992; Powell et al., 1992). Germline mutations in *APC* also form the genetic basis of familial adenomatous polyposis (FAP), an inherited form of the disease that is characterized by hundreds of colorectal polyps that progress to cancerous lesions if left untreated (Leoz et al., 2015). This makes a comprehensive understanding of the normal interactions and functions of APC crucial for effectively targeting *APC* mutant cells.

The tumor suppressive function of APC has been mainly attributed to its role in Wnt signaling. In conjunction with Axin, APC acts as a scaffold for the β-catenin destruction complex, thereby limiting the transcription of pro-proliferative β-catenin target genes in the absence of Wnt ligands (Stamos and Weis, 2013). The vast majority of *APC* mutations result in the translation of a truncated protein and consequent deregulation of Wnt signaling (Miyoshi et al., 1992; Powell et al., 1992). Nevertheless, Wnt-independent roles of APC also contribute to its function as a tumor suppressor. This is exemplified by the rare detection of mutations in other Wnt signaling components, including β-catenin, in CRC (Polakis, 2000; Polakis, 2007). Although deletion of Apc in the intestinal epithelium in mice phenocopies homozygous truncation mutations, it leads to more rapid onset of tumors despite lower levels of Wnt activation (Cheung et al., 2010). It thus emerges, that loss of wild-type (WT) APC confers additional advantages to cells beyond β-catenin-mediated proliferation, and that these advantages appear to be particularly important for intestinal tumorigenesis.

Because the intestinal epithelium is constantly renewed, cells with potentially oncogenic mutations are normally quickly eliminated from the tissue (Näthke, 2004). Notably, expression of truncated APC inhibits cell migration *in vitro* and *in vivo* and induces defects in microtubule networks required for directed cell migration (Mahmoud et al., 1997; Etienne-Manneville and Hall, 2003; Etienne-Manneville et al., 2005). The resulting prolonged residence of *APC*-mutant cells in the gut epithelium could allow these cells to acquire additional mutations that confer growth advantages to permit tumors to form (Näthke, 2004; Wong et al., 1996). Importantly, intestinal microadenomas do not display increased proliferation, indicating that defects in migration, rather than proliferation, are the initiating mechanism of polyp formation (Oshima et al., 1997).

A variety of proteins have been described to interact with APC in addition to β-catenin destruction complex components (Nelson and Näthke, 2013). However, proteome-wide studies of APC-binding proteins are limited to interactome and yeast-two-hybrid experiments with overexpressed, tagged and/or fragments of APC (e.g. Breitman et al., 2008; Brandyopadhyay et al., 2010; Hein et al., 2015; Song et al., 2012). Using tagged APC in interaction studies is problematic because the C-terminal PDZ-binding domain must remain free to interact with other proteins (Harris and Lim, 2001). Similarly, the N-terminal oligomerization domains rely on coiled-coil formation and may be compromised by N-terminal tags (Joslyn et al., 1993). To overcome these limitations, we used label-free affinity-enrichment mass spectrometry (AE-MS) to identify a more comprehensive set of interacting partners of endogenous, non-tagged APC. Furthermore, we applied an untargeted global approach using tandem mass tag (TMT)- based and label-free MS to identify proteins that are – similarly to, but independently of β-catenin – regulated by APC in their abundance (Figure 1A).

**Figure 1.**
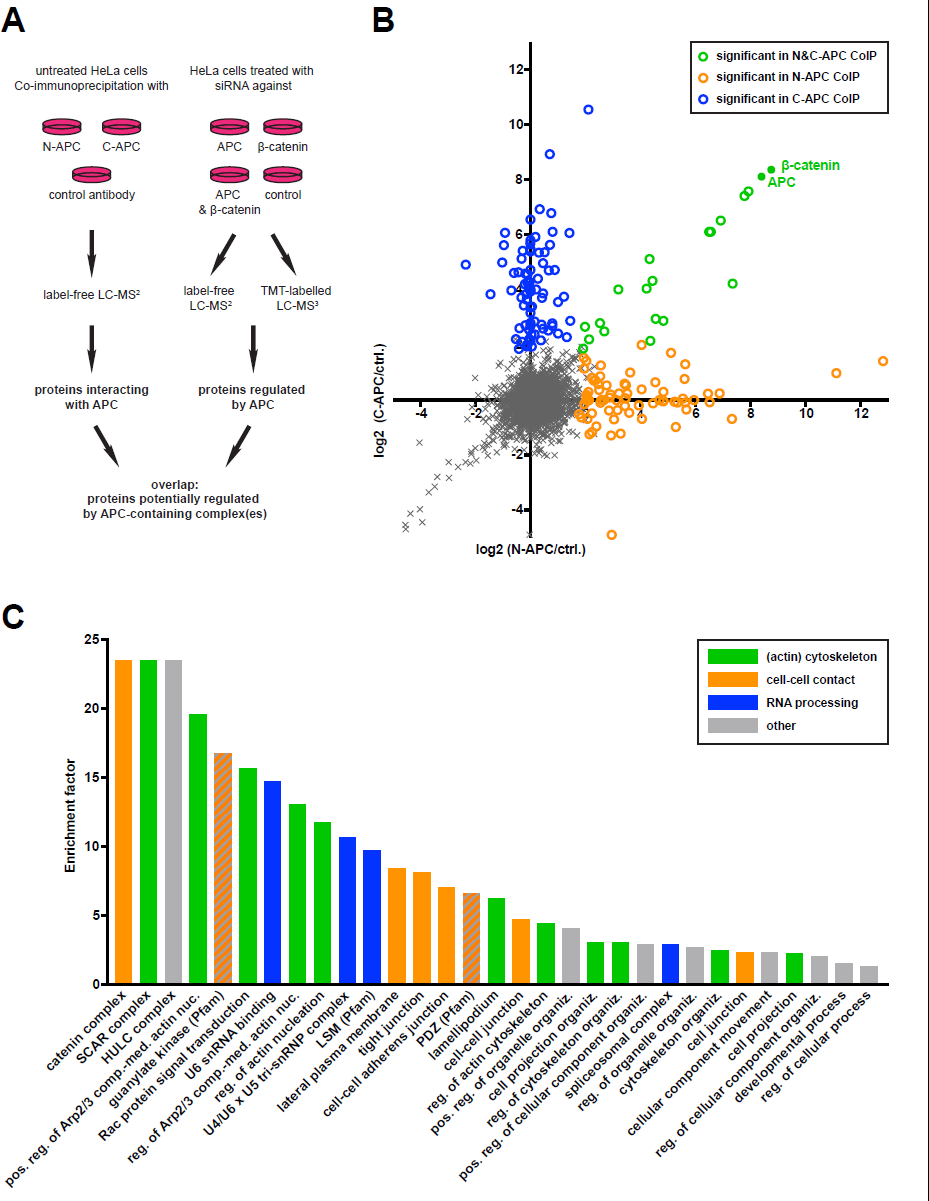
Identification of APC-interacting and Eregulated proteins. A Experimental Outline. Proteins in APC-containing complexes and changes in protein expression in response to siRNA-mediated depletion of APC and/or β-catenin were analyzed by label-free and TMT-based mass spectrometry. The overlap between the two data sets constitutes potential targets of alternative APC-containing (destruction) complexes. **B** Proteins significantly enriched in C- and/or N-APC Co-IPs. Log2 fold change in mean LFQ intensities between N-APC Co-IP vs. control IP (*x*-axis) plotted against log2 fold change in mean LFQ intensities between C-APC Co-IP vs. control IP (*y*-axis). Significance determined by two-sided t-test with permutation-based FDR <0.01 and s0 = 2 used for truncation (Tusher, 2001). See also Figure S1, S2 and Table S1. **C** GO, Pfam and KEGG terms significantly enriched in the APC interactome data set. Enrichment calculated by Fisher Exact Test, significance determined by BenjaminiHochberg corrected FDR <0.02. Abbreviations: pos. - positive, reg. - regulation, comp.-med. - complex-mediated, nuc. - nucleation, organiz. - organization. See also Table S2.

## Results

### Identification of APC-interacting proteins by affinity enrichment-mass spectrometry (AEMS)

For our initial discovery experiments, we used HeLa cells, which express relatively high amounts of WT APC that can be efficiently depleted by siRNA. This allowed us to measure both, protein binding to and regulation by APC, in the same cell line. APC-containing protein complexes were co-immunoprecipitated using two APC-specific monoclonal antibodies (ABs) that recognize N- and C-terminal domains, respectively. An isotype-matched AB against the viral V5 peptide was used as control. Coimmunoprecipitation (Co-IP) with each AB was performed in quadruplicate. Following ingel tryptic digestion and peptide extraction, samples were analyzed by label-free tandem mass spectrometry (LC-MS/MS). Of 5,521 measured proteins, 4,016 were detected in all four replicates of Co-IP’s with either AB and only these were considered for further analysis. Pearson correlation coefficients >0.9 for label-free quantification (LFQ) intensities measured across replicates and a clear separation of N-APC, C-APC and control Co-IPs by principal component analysis (PCA) indicated good experimental reproducibility (Figures S1A and S1B). Significant enrichment of proteins in APC-specific versus control Co-IPs was determined by considering both permutation-based FDR (<0.01) and LFQ intensity fold-change (Figure S1C).

In total, 171 proteins were significantly enriched in APC-specific Co-IPs (Figure 1B and Table S1). These proteins will be referred to hereafter as ‘APC interactome’. Eighty and 71 proteins were exclusively enriched in either C-APC or N-APC Co-IPs, respectively. AB binding to APC is likely affected by protein interactions at domains close to or overlapping with the AB epitopes. This could explain co-immunoprecipitation of distinct interactors with different APC-specific ABs. Consistently, C-APC and N-APC ABs immunoprecipitated overlapping, but distinct, pools of APC protein that may contain different subsets of binding partners (Figure S1D). Twenty proteins, including APC itself, were significantly enriched in both APC Co-IPs and only half of these were previously described APC interactors (Orchard et al., 2014; Stark, et al., 2006; Table S1).

### Validation of APC AE-MS findings in CRC cells

The majority of potential APC interaction partners identified here have not been reported previously. To rule out a HeLa cell-specific enrichment of proteins in APC CoIPs, we validated our AE-MS results in the human colon carcinoma cell line HCT116-Haβ92. These cells express WT APC and are hemizygous for WT β-catenin (the mutant allele present in the parental cell line has been deleted; Kim et al., 2002). Thirteen of the novel APC-interacting proteins were selected to cover the range of biological functions represented in the data set and antibody availability. APC Co-IPs were performed using HeLa and HCT116-Haβ92 cell lysate and co-immunoprecipitated proteins were detected by western blotting (WB, Figure S2). Consistent with results obtained by AEMS, all 13 proteins were enriched in both cell lines. One exception was GIT2, which could not be detected in C-APC Co-IPs from HeLa cell lysate by WB.

### The APC interactome is enriched for epithelial-specific GO cellular component terms

To identify underlying functional patterns, we analyzed the enrichment of gene ontology (GO), protein family (Pfam), and Kyoto encyclopedia of genes and genomes (KEGG) terms in the APC interactome (compared to all measured proteins) by Fisher Exact Test. Thirty-one terms were significantly over-represented (Benjamini-Hochberg FDR <0.02); the majority can be broadly categorized into three cellular processes: (actin) cytoskeleton organization, cell-cell contact establishment, and RNA processing (Figure 1C and Table S2). APC-interacting proteins associated with cytoskeletal organization included the known binding partners Disks large homolog 1 (Dlg1) and Rho GTPase-activating protein 21, and newly identified interactors, including several SCAR complex components. SCAR/WAVE proteins promote the nucleation of actin filaments by regulating the Arp2/3 complex (Pollitt and Insall, 2009). The enrichment of terms linked to RNA processing is consistent with APC’s role as an RNA-binding protein (Preitner, et al., 2014). Strikingly, many of the enriched terms are associated with cell-cell contacts and constitute components characteristic of epithelial cells, for example “lateral plasma membrane”, “tight junctions”, “cell-cell adherens junction”, and “guanylate kinase”. All proteins present in the latter category are members of the membrane associated guanylate kinase (MAGUK) family that is implicated in the establishment of epithelial cell polarity (Caruana, 2002). APC contains a PDZ-binding domain and, indeed, our data set was enriched for PDZ domain-containing proteins. These included several previously unknown interactors, such as Rap guanine nucleotide exchange factor 2 and 6. In summary, the results of the enrichment analysis were consistent with known functions of APC and provide evidence for the importance of APC specifically in epithelial cells.

### Generation of an integrated APC interaction network

Our experiments identified many new interaction partners that could facilitate and coordinate known functions and/or link APC to processes not previously considered. To understand the relationship between interactions partners and known and potential new APC functions, we tested how our interactome integrated into a network of previously identified APC-binding proteins. Our interactome data set overlaid well with and added substantially to the network of known APC-binding proteins (Figure 2). Most proteins common to both data sets belonged to a cluster of β-catenin destruction complex components. In addition, the integrated network revealed many direct and indirect high-confidence links between newly identified and known APC interactors suggesting potential APC-interacting protein complexes. In addition to the ‘β-catenin destruction complex’ cluster, several sub-networks emerged from this analysis. Two of these included proteins associated with ‘LSM protein family’ and ‘SCAR complex’, respectively, and both categories were enriched in our APC interactome data set (Figure 1C). Members of the Striatin-interacting phosphatase and kinase (STRIPAK) complex and cytoskeleton elements – including actin and microtubule-binding proteins and intermediate filament components – formed two additional sub-networks.

**Figure 2.**
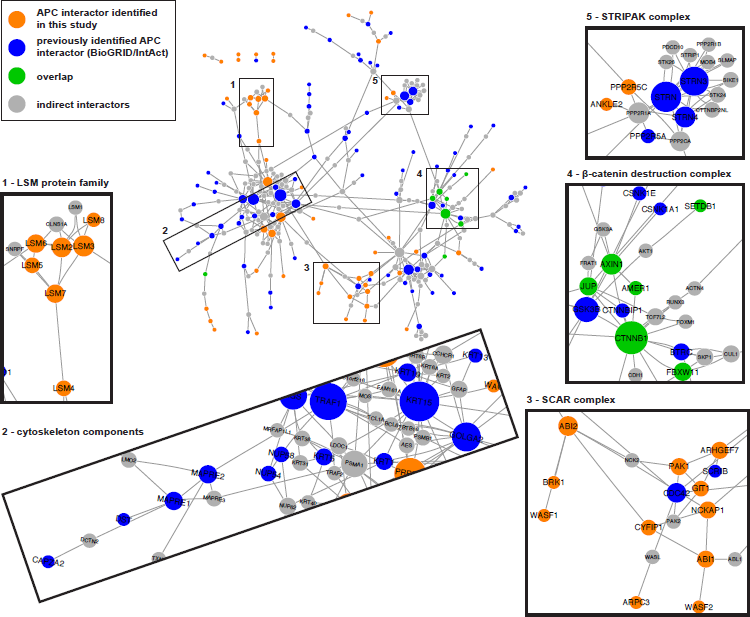
APC interactome network. Network integrating known (blue), newly identified (orange), and indirect (grey) APC interaction partners. Nodes are labelled with corresponding gene names and node size correlates with degree of connectivity, i.e. number of edges. Components of distinct protein complexes (1, 3-5) and proteins associated with the cytoskeleton (2) cluster together in sub-networks. See also Figure S3 and Network S1.

To validate our network analysis, we generated a control network using 171 proteins randomly selected from the APC AE-MS data set (Figure S3). Compared to the random selection, our interactome exhibited superior integration into the network of known APC-binding partners as indicated by the higher number of proteins connected to the main network and more connections, including direct links to known interactors. In summary, the integrated APC interaction network revealed distinct classes of APC- binding proteins and highlighted novel components of potential APC-interacting protein complexes.

### APC affects the abundance of many proteins independently of β-catenin

The identified APC interactome adds to a growing list of binding partners that appear unrelated to Wnt signaling components (Nelson and Näthke, 2013). We therefore aimed to investigate whether APC is involved in the regulation of proteins other than β-catenin, and independently of β-catenin-mediated transcription. To this end, we depleted APC alone or together with β-catenin from HeLa cells using siRNA and measured changes in protein abundance by mass spectrometry (MS). Cells were harvested 72 hours after transfection and efficient knockdown was confirmed by WB (Figure S4A). Simultaneous knockdown of APC and β-catenin abrogated β-catenin target gene activation, as verified by the inhibition of *AXIN2* mRNA transcription compared to APC siRNA-treated cells (Figure S4B). For each siRNA combination, we analyzed four and two experimental replicates by label-free and TMT MS, respectively. Downstream analysis was applied to 6,923 proteins measured in all eight samples by TMT MS and 4,927 proteins measured in at least three replicates of at least one condition by label-free MS (Tables S3 and S4). Reproducibility between replicates was very good, as indicated by Pearson correlation coefficients >0.97 and a clear separation of distinct siRNA treatments by PCA (Figure S4C and D).

To identify proteins that changed in abundance in response to APC depletion, but independently of β-catenin, we compared TMT/LFQ intensities across conditions of all measured proteins to an “ideal” intensity profile of a hypothetical β-catenin-independent APC target (defined in legend to and indicated in red in Figure 3A and S5A). The 200 proteins with profiles most similar to the ideal negative and positive APC target, respectively, were selected based on Pearson correlation. Significant β-catenin-independent APC targets were determined by applying an additional cut-off of >1.5 fold-change in APC and APC+β-catenin siRNA-treated samples relative to control with a *q-*value <0.05 (TMT)/0.1 (LFQ). By TMT MS we identified 53 and 85 proteins that significantly increased and decreased, respectively, in response to APC depletion in a β-catenin-independent manner; by LFQ MS 11 proteins increased and 11 decreased (Figure 3A/B and S5A/B). Four negatively and seven positively regulated proteins were common to both data sets. This group of proteins was not enriched in distinct GO terms (data not shown), but spanned a range of cellular functions including apoptosis, ion transport, actin organization, and proliferation.

**Figure 3.**
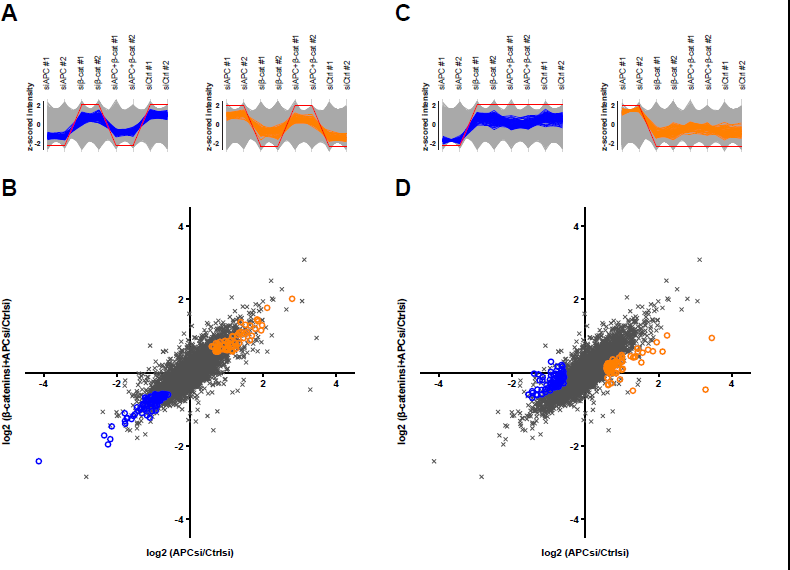
β-catenin-dependent and – independent APC targets identified by TMT MS. Profiles of z-scored TMT intensities of all measured proteins across samples. Protein identified as negative and positive β-catenin-independent APC targets are shown in orange and blue, respectively. Red lines show profiles for hypothetical ‘ideal’ targets that increase/decrease in response to APC depletion, but irrespective of a change in βcatenin. **B** Log2 fold change in mean TMT intensities between APC siRNA and control siRNA treated samples (*x*-axis) plotted against the log2 fold change in mean TMT intensities between β-catenin+APC siRNA and control siRNA treated samples (*y*-axis). Proteins selected based on their TMT intensity profiles in **A** are shown in orange and blue, respectively. **C** and **D** Same as A, but for β-catenin-dependent APC targets. See also Figure S4 and S5.

To compare APC’s β-catenin-dependent and – independent effects on protein expression, we also identified proteins that changed in abundance in response to APC depletion in a β-catenin-dependent manner. The number of these proteins was similar to those regulated independently of β-catenin: 64 and 37 were negatively regulated, 86 and 103 were positively regulated when detected by TMT and LFQ MS respectively (Figure 3C/D and S5C/D, Table S3 and S4). Collectively, our results indicate that APC plays a role in the regulation of many proteins that are not β-catenin targets.

### Some β-catenin-independent APC targets are also deregulated in human cancer

To determine if any of the identified β-catenin-independent APC targets are implicated in CRC, we compared our results with a dataset describing proteomic changes in human colorectal adenoma and adenocarcinoma (Wiśniewski, et al., 2015). Nineteen proteins present in our APC target list were also found to be dysregulated – in the same direction – in human adenomas and/or carcinomas, i.e. they increased/decreased in response to APC depletion independently of β-catenin and were up-/down-regulated in diseased compared to healthy tissue, respectively (Table 1). These results highlight that mis-expression of some proteins in CRC could be a direct consequence of loss of WT APC rather than deregulated Wnt signaling.

**Table 1.**
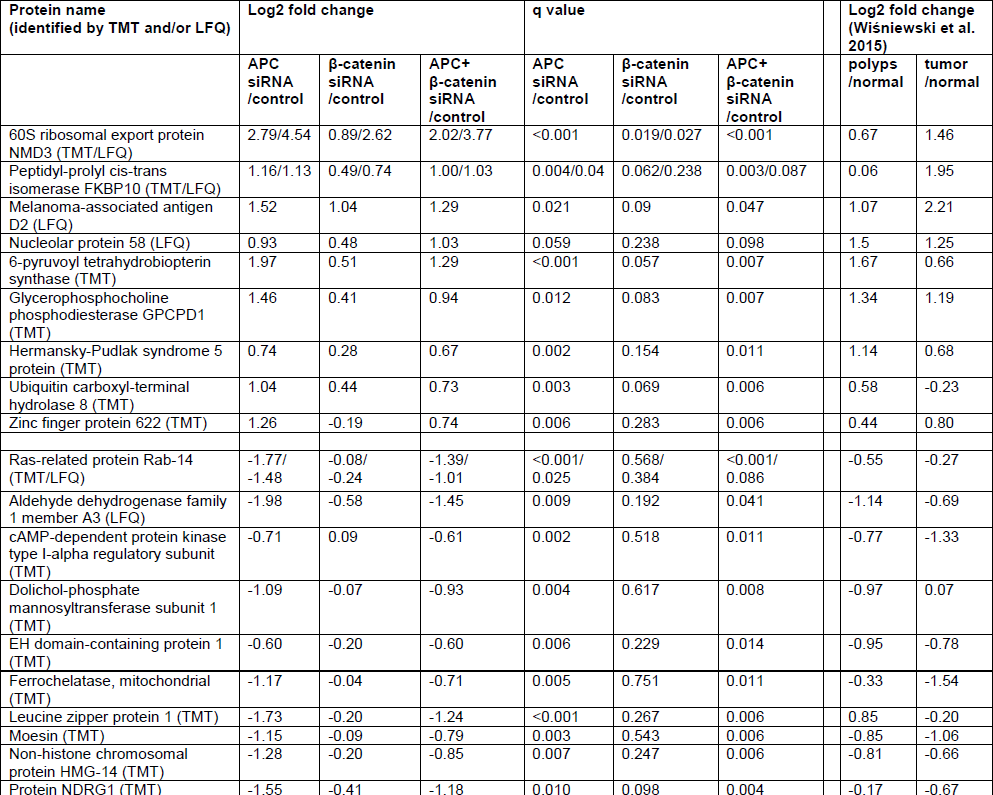
β-catenin-independent APC targets deregulated in CRC. See also Table S3 and S4. Overlap between β-catenin-independent APC targets identified in this study and proteins mis-expressed in colorectal polyps and/or tumors (dataset by Wiśniewski et al., 2015). The experiment (TMT-based (TMT) or label-free (LFQ) mass spectrometry) in which the protein was identified as a significant β-catenin-independent APC target is indicated in brackets after the protein name.

### Misshapen-like kinase 1 (MINK1) is a potential target of an alternative APC-containing destruction complex

Our data provided an opportunity to better understand how APC can regulate the abundance of proteins other than β-catenin. Proteins that are regulated by APC-containing degradation or stabilization complexes are expected to interact with APC. From the group of β-catenin-independent APC targets identified by total proteomics analysis, six fulfilled this condition. Amongst these, MINK1, which accumulated in response to APC depletion, stood out as a potentially druggable serine/threonine kinase and thus worth pursuing further. Little is known about the functions of MINK1 or the regulation of its activity and/or its abundance. Existing data implicate MINK1 in cell adhesion, cell migration and planar cell polarity (PCP; Hu et al., 2004; Daulat et al., 2012; Mikrykov and Moss, 2012) - processes crucial for epithelial biology. Furthermore, MINK1 kinase activity is required for completion of cytokinesis (Hyodo et al., 2012). Importantly, these processes are also deregulated in *APC* mutant tissues (Wong et al., 1996; Mahmoud et al., 1997; Caldwell et al., 2007). Moreover, TRAF2 and NCK interacting protein kinase (TNIK), which shares high sequence homology with MINK1, is emerging as a promising target for CRC therapy, as it regulates the activity of the TCF-4/β-catenin transcription complex (Yamada and Masuda, 2017; Masuda, et al., 2016).

### MINK1 interacts with full length and truncated APC

We validated the interaction between MINK1 and full-length APC by Co-IP and WB in two cell lines (Figure 4A). In agreement with results obtained by MS, MINK1 was only enriched in Co-IPs with N-APC AB (data not shown). To rule out that coimmunoprecipitation of MINK1 in N-APC IPs was caused by AB cross-reactivity, we repeated the experiment with lysate from APC siRNA-treated cells. Confirming its specific enrichment in APC protein complexes, the amount of co-immunoprecipitated MINK1 correlated with the levels of APC present in IP lysates (Figure S6A). We next tested whether MINK1 could also interact with truncated APC expressed in CRC cells. MINK1 co-immunoprecipitated with APC fragments in both SW480 and Colo320 cells, which express ˜220 kDa and ˜90 kDa N-terminal APC fragments, respectively (Figure 4B). These results validate MINK1 as a novel APC interaction partner and suggest that the interaction between the two proteins is mediated by domains in the N-terminal third of APC.

**Figure 4.**
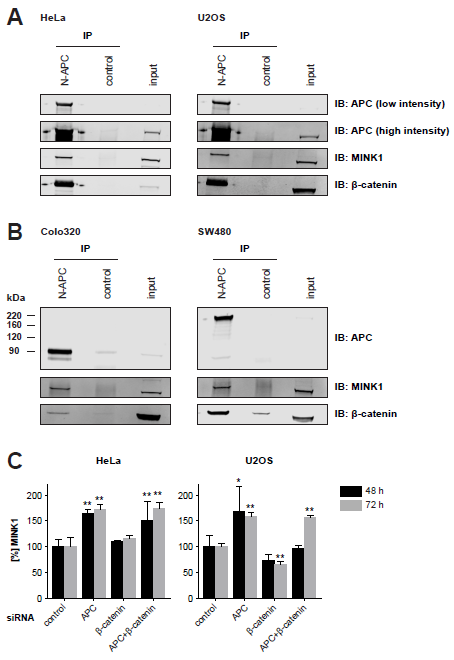
MINK1 binds to and is negatively regulated by APC. **A** Co-IP of MINK1 with full-length, endogenous APC in HeLa and U2OS cells. See also Figure S6A. **B** Co-IP of MINK1 with C-terminally truncated APC in Colo320 (APC mutation in codon 811) and SW480 (APC mutation in codon 1338) CRC cells; both cell lines lack the second WT allele (full-length APC migrates at approximately 300 kDa). βcatenin was only significantly enriched in APC Co-IPs in SW480 cells, which - in contrast to Colo320 cells - express an APC fragment that retains some of its β-catenin binding sites. **C** Changes in MINK1 proteins levels in response to siRNA-mediated depletion of APC and/or β-catenin measured by WB. Shown are means and SD relative to control samples from four independent transfections. Significance relative to control determined by two-way ANOVA followed by Dunnett’s multiple comparison test; *: p value < 0.05, **: p value < 0.01. See also Figure S6B and C.

### MINK1 is negatively regulated by APC independently of β-catenin

Consistent with our proteomics data, MINK1 levels measured by WB significantly increased above 1.5-fold in response to APC depletion in HeLa and U2OS cells and this accumulation was independent of changes in β-catenin (Figure 4C and S6B).

Transfection with β-catenin siRNA alone did not have a consistent effect on MINK1 levels. To exclude potential siRNA off-target effects, HeLa cells were transfected with individual APC siRNAs (APC siRNA #1 - #3) and a combination of these (APC siRNA pool), respectively. Similar to results obtained with the siRNA pool, transfection with either of the individual siRNAs efficiently decreased APC levels and produced a concomitant increase in MINK1 protein (Figure S6C). We validated the effect of APC loss on MINK1 levels *in vivo* by measuring protein expression in intestinal tissue from *Apc* mutant and WT mice. Mink1 protein was increased by 2.3-fold (±0.4 SD) in *Apc^Min/+^* versus control animals (n=4 per genotype, Figure 5A and 5B). In addition, we addressed whether this regulatory relationship is conserved across species. Misshapen (Msn), the closest orthologue in *Drosophila melanogaster*, is, implicated in PCP, cell migration, and adhesion, similar to MINK1. These processes are important in follicular epithelial cells for shaping the *Drosophila* egg chamber during oogenesis (Paricio et al., 1999; Horne-Badovinac et al., 2012; Lewellyn et al., 2013). Two partially redundant paralogues of APC (APC1 and APC2) are present in *Drosophila* (Akong et al., 2002a; Akong et al., 2002b) and we induced mitotic recombination to generate mosaic follicular epithelia in egg chambers carrying clones of double *APC1*, *APC2* mutant cells (marked by loss of NLS::RFP expression). We then measured levels of a YFP-fused Msn (Msn::YFP) protein expressed from its endogenous locus (Lowe et al., 2014; Lye et al., 2014) using live microscopy. Direct comparison of mutant and control neighboring cells revealed that Msn::YFP levels were 28±12% higher in cells that did not express APC1 and APC2 (n=10 clones from 10 different egg chambers, Figure 5C-E). Together, our results demonstrate that APC limits the abundance of MINK1 independently of β-catenin. Correspondingly, loss of WT APC resulted in increased levels of MINK1 *in vitro* and *in vivo*.

**Figure 5.**
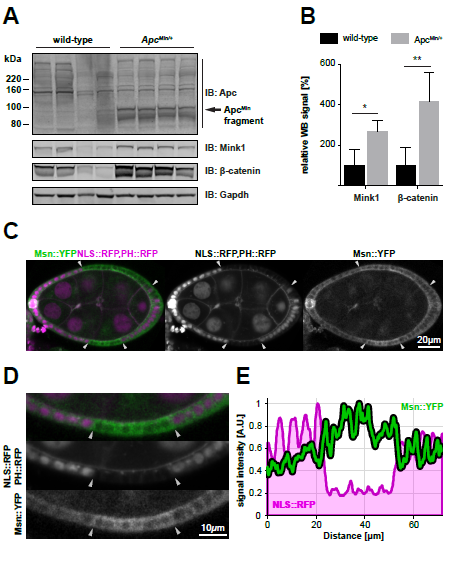
Mink1/Msn levels increase in response to Apc loss *in vivo*. **A** Expression of Mink1 in small intestinal tissue lysate from WT and Apc^Min/+^ mice measured by WB. The Apc^Min^ fragment of approximately 90 kDa was present in mutant mice, but full-length Apc (˜310 kDa) was not detectable. **B** Quantification of WB shown in A. Shown is the relative mean WB signal normalized to Gapdh. Significance relative to WT samples determined by un-paired, two-tailed t test; p value: * < 0.05, ** < 0.01. **C** Live stage 8 *Drosophila* egg expressing NLS::RFP and PH::RFP (magenta) under the control of a ubiquitous promoter, and endogenous Msn::YFP (green). Two large *APC1*, *APC2* double mutant clones within the follicular epithelium are identified by the absence of NLS::RFP and delimited by arrowheads. **D** Magnification of one *APC1*, *APC2* double mutant clone displayed in C. **E** Intensity profiles of RFP and Msn::YFP signal along the follicular epithelium.

### Parallels between the regulation of MINK1 and β-catenin protein abundance

Our results suggest that MINK1 is a substrate of an alternative APC-containing destruction complex. We hypothesized that APC regulates the abundance of MINK1 – similarly to β-catenin – post-transcriptionally. Transfection with APC siRNA resulted in a significant up-regulation of *AXIN2* mRNA (up to 50-fold after 72 h), and this increase was efficiently inhibited when APC and β-catenin were depleted simultaneously. In contrast, *MINK1* mRNA increased moderately (up to 2.3-fold) but changes in *MINK1* mRNA did not correlate with changes in MINK1 protein abundance and were inconsistent with specific knockdown of APC and/or β-catenin (Figure 6A and S7A, Figure 4C). Based on the role of APC in the phosphorylation and ubiquitination of β-catenin, we investigated changes in post-translational modifications (PTMs) of MINK1 as a readout for its regulation by APC. PTMs on MINK1, which was immunoprecipitated from cells treated with control or APC siRNA, respectively, were analyzed by label-free MS. We detected 34 post-translationally modified sites; the majority was unaffected by APC depletion (Table S5). In contrast, phosphorylation at serine 674 was significantly reduced in APC knockdown versus control samples (Figure 6C and S7C). S674 lies in the central region between the N-terminal kinase domain (aa 25-289) and the C-terminal regulatory CNH domain (aa 1014-1312) of MINK1. This residue is a predicted target site for GSK3 and CK1 (The Eukaryotic Linear Motif (ELM) resource; Dinkel et al., 2016) and both cooperate with APC in the β-catenin destruction complex.

**Figure 6.**
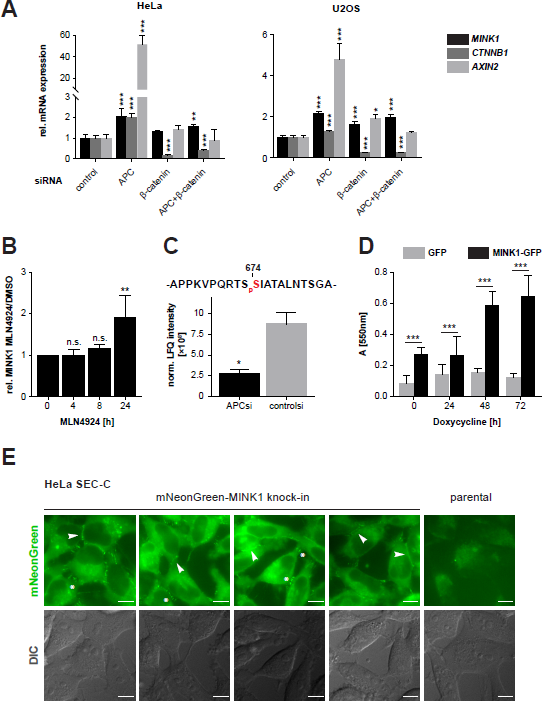
MINK1 localizes to cell-cell junctions and its overexpression enhances cell adhesion. See also Figure S7. **A** mRNA expression of *MINK1*, *CTNNB1*, and *AXIN2* measured by RT-qPCR 72 h after siRNA transfection. Indicated are mean expression levels relative to *ACTB* expression with SD from four independent transfections. Significance determined by one-way ANOVA followed by Dunnett’s multiple comparison test; p value: * < 0.05, ** < 0.01, *** < 0.001. Note, the same HeLa *AXIN2* mRNA quantification data is also shown in Figure S4B. See also Figure S7A. **B** MINK1 protein levels in HeLa cells after treatment with neddylation inhibitor MLN4924 [3 μM] as measured by WB. Shown are relative mean signals normalized to DMSO-treated samples with SD from three independent experiments. Inhibition of E3 ubiquitin ligase activity by MLN4924 resulted in an approximately two-fold increase of MINK1 protein after 24 h. Significance determined by one-way ANOVA, **: p value < 0.01. See also Figure S7B. **C** Phosphorylation of MINK1 at S674 72 h after transfection with APC and control siRNA, respectively, in HeLa cells as measured by MS. Shown are mean LFQ intensity values normalized to total MINK1 protein with SD from four independent transfections. Significance determined by Benjamini-Hochberg corrected FDR <0.05, *: q value = 0.047. See also Figure S7C and Table S5. **D** Adhesion assay with U2OS cells overexpressing MINK1-GFP and GFP, respectively. Adhesion to collagen matrix after one hour was quantified by staining of firmly attached cells with Crystal Violet. Indicated is mean absorbance with SD of independent experiments with two different GFP/MINK1-GFP clones and eight technical replicates/condition. Significance determined by two-way ANOVA followed by Sidak’s multiple comparison test; p value: *** < 0.0003. See also Figure S7D. **E** Live imaging of HeLa SEC-C cells expressing endogenously mNeonGreen-tagged MINK1. Enrichment of mNeonGreen signal is visible at lateral plasma membranes in areas of cell-cell contact (arrow heads) and tips of protrusions contacting neighboring cells (asterisks), but is absent from contact-free plasma membrane areas. Scale bars: 10 μm. See also Figure S7F.

We further tested whether the degradation of MINK1, similarly to β-catenin, was dependent on the action of an E3 ubiquitin ligase. Treatment with the NEDD8-activating enzyme selective inhibitor MLN4924, which inhibits cullin-RING ubiquitin ligase activity (Soucy, et al., 2009), reproducibly induced a two-fold increase in MINK1 after 24 h (Figure 6B and S7B). This indicates that MINK1 potentially serves as a substrate for the cullin-RING ubiquitin ligase/proteasome machinery. Collectively, our results reveal many similarities between the regulation of MINK1 and β-catenin protein abundance suggesting that APC regulates MINK1 in a process analogous to its role in the β-catenin destruction complex.

### MINK1 localizes to cell-cell junctions and enhances cell adhesion

To address how elevated MINK1 could contribute to cellular processes affected by *APC* mutations, we determined its sub-cellular localization. Immunofluorescence (IF) staining using an anti-MINK1 AB in U2OS cells showed an enrichment of signal in the perinuclear region (Figure S7E). This is in agreement with a previous study describing an overlap between the localization of MINK1 and the Golgi network (Hu et al., 2004). To confirm specificity of the AB, we generated MINK1 knockout cell lines using CRISPR-Cas9 (Figure S7F). The perinuclear IF signal persisted in these cells, suggesting cross-reactivity of the MINK1 AB with Golgi components (Figure S7E). As shown by WB, the MINK1 AB used for IF indeed recognized additional proteins of similar molecular weights to MINK1 (Figure S7F, the MINK1 AB used for WB in this study was unsuitable for IF).

We employed CRISPR-Cas9 to generate cells expressing endogenously mNeonGreen-tagged MINK1, enabling us to study its localization live in un-fixed cells (Figure S7F). Although fluorescence intensity was low, mNeonGreen signal was clearly enriched at tips of protrusions and at lateral plasma membranes in areas of cell-cell contact (Figure 6E). Conversely, no signal enrichment was detected in ‘free’ regions of the plasma membrane without adjoining cells.

Guided by these findings, we analyzed the effects of elevated MINK1 on cell adhesion. We generated stable U2OS cell lines with doxycycline-inducible expression of MINK1-GFP or GFP alone (Figure S7D) and measured cell adhesion. Overexpression of MINK1-GFP resulted in a significantly higher number of cells attached to collagen compared to controls, and an increase in cell attachment over time (Figure 6D). Based on the localization of MINK1 to cell-cell contacts, we propose that its overexpression enhances cell adhesion which leads to the formation of larger groups of cells that then cooperatively attach to the matrix.

## Discussion

It is becoming increasingly clear that Wnt-independent roles contribute to the function of APC as a tumor suppressor. We aimed to elucidate – on a global scale – the diverse molecular roles of APC, with an emphasis on its functions beyond the β-catenin destruction complex. To this end, we applied an untargeted approach using label-free and TMT-based MS to assemble an APC interactome and, furthermore, to identify the β-catenin-independent APC-regulated proteome. These data sets provide a useful resource for the identification of proteins that participate in and coordinate Wnt-independent functions of APC.

Using APC AE-MS, we identified 170 potential APC interactors, of which only twelve were identified previously (Figure 1B). In contrast to previous interaction studies, we used endogenous, full-length, and non-tagged APC. The identification of additional PDZ domain-containing APC interaction partners highlighted the benefit of this approach. The emergence of distinct functional clusters related to β-catenin destruction, cytoskeleton regulation, RNA processing, etc. within the generated APC interaction network further confirmed APC as a multi-functional protein (Figure 2). Strikingly, the APC interactome was highly enriched for proteins that are part of cellular components characteristic for epithelial cells, as well as MAGUK protein family members, and PDZ-domain containing proteins (Figure 1C). MAGUK proteins are implicated in the establishment of epithelial cell polarity (Caruana et al., 2002). Furthermore, the function of APC in epithelia is – at least partly – mediated by PDZ domain-containing proteins (Mimori-Kiyosue et al., 2007). In addition, STRIPAK complex components formed a highly connected cluster within the APC interaction network (Figure 2). APC and the STRIPAK component Striatin localize interdependently to cell-cell junctions in epithelial cells and depletion of Striatin and APC affects tight junction organization (Breitman et al., 2008). It is conceivable that binding to APC regulates the sub-cellular localization, activity, and/or expression of these epithelial-characteristic proteins, in turn controlling cellular adhesion and establishment of epithelial polarity. Reduced expression of proteins involved in cell adhesion in *APC* mutant cells supports this hypothesis (Halvey et al., 2012). Some of these binding partners might affect APC itself, e.g. by controlling its localization, similarly to AMER1 (Grohmann et al., 2007). The enrichment of epithelial-specific proteins raises the possibility that the APC interactome differs between epithelial and non-epithelial cells. Measuring such differences will provide useful insights into the mechanisms that regulate APC function in different tissues and further improve our understanding of the phenotypes associated with APC loss. Such studies could reveal why *APC* germ line mutations in FAP patients result in cancerous lesions of the gut epithelium, while other organs often remain unaffected.

In addition to its established role in the regulation of microtubules, APC can also bind and bundle actin filaments via its basic domain *in vitro* (Moseley et al., 2007) and nucleate actin assembly in cooperation with mDia (Okada et al., 2010). Furthermore, APC impacts on the activity of the actin regulators Rac and Cdc42 through interactions with Asef and the Rac/Cdc42-effector IQGAP1 (Kawasaki et al., 2000; Watanabe et al., 2004). In addition to the known interactor Rho GAP21, we identified four Rho/Rap GEFs (Rap GEF2 & 6, Rho/Rac GEF FGD1, Rho GEF7) as novel APC binding proteins. Together with the identification of SCAR/WAVE complex components as APC interaction partners, this provides further evidence for the involvement of APC in the regulation of actin dynamics. Moreover, GIT1 and 2, which serve as GAPs for ARF6 and are involved in vesicle transport (Donaldson and Jackson, 2011), were found to interact with APC. Links between APC and vesicular transport have not yet been explored but may be responsible for the differential expression in plasma membrane integral proteins, such as ion channels and transporters, between healthy and cancerous colon tissue (Wiśniewski et. al, 2015).

Measuring proteome-wide effects of APC loss revealed a set of β-catenin-independent APC targets, supporting a role of APC in the regulation of protein abundance beyond the β-catenin destruction complex (Figure 3 and S5). Similar to the effect on β-catenin, depletion of APC resulted in the accumulation of some proteins, while the levels of others were negatively affected, suggesting that APC can also inhibit degradation of some its targets. Strikingly, the number of proteins regulated by APC independent and dependent of β-catenin is very similar. It is important to acknowledge that untargeted MS is biased towards detection of more abundant proteins.

Consistently, many established, but low-abundant, β-catenin targets, such as Myc and Axin2, were not detected and are thus absent from our analysis. However, this bias operates in both sets of targets equally. Changes in the abundance of individual APC targets could result from alterations in PTMs and/or protein stability – as is the case for β-catenin. This is supported by previous findings in *Drosophila*, where loss of APC2 causes proteome-wide and β-catenin-independent changes in post-translational modification that also affect protein stability of some proteins (Blundon et al., 2016). In addition, effects on transcription may also contribute to the observed differences in protein abundance.

At present, it remains unclear how changes in these APC targets contribute to the functional consequences of APC loss observed *in vivo*. As a first step towards addressing this question, we compared our data set with data describing proteome-wide changes in colon adenomas and carcinomas (Wiśniewski et. al, 2015). Several of the β-catenin-independent APC targets we identified were also found to be deregulated in colorectal adenomas and/or tumors (Table 1). Among these, NDRG1, which was downregulated in APC-depleted cells in our study and also in cancerous tissue, might be of particular interest. NDRG1 has been established as a tumor suppressor in CRC cells based on its negative effects on metastasis and apoptosis (Fang et al., 2014; Mi et al., 2017).

Collectively, our results suggest that part of the protein expression changes observed in CRC are independent of increased Wnt target gene expression. Investigating the functional impact of these changes will further help to elucidate how APC loss contributes to cancer development beyond de-regulated β-catenin. Accounting for these effects will be especially important when considering cancer therapy, as they reveal that consequences of mutant APC protein cannot be fully rectified by restoring normal Wnt signaling.

From the group of six proteins we found bound to and regulated in their abundance by APC independently of β-catenin, MINK1 stood out as it is a potentially druggable serine/threonine kinase. MINK1, also known as MAP4K6, activates the JNK and p38 MAPK signaling pathway (Dan et al., 2000; Hu et al., 2004; Chuang et al., 2016; Lim et al., 2003) and has been suggested to function in parallel to MST-1 and E2 in the Hippo pathway (Meng et al., 2015). De-phosphorylation of MINK1 by PP2A reduces its kinase activity (Hyodo et al., 2012), but additional upstream regulators of MINK1 activity and/or expression are largely unknown. Our data suggested that APC might be directly involved in regulating the abundance of MINK1.

We validated the interaction between MINK1 and APC, as well as its negative and β-catenin-independent regulation by APC, in two cell lines (Figure 4). RT-qPCR analysis indicated that APC regulates MINK1 predominantly post-transcriptionally (Figure 6A and S7A). This was further supported by our MINK1 PTM analysis, suggesting that APC facilitates the phosphorylation of MINK1 at S674, which may be a signal for its degradation (Figure 6C). Together with the accumulation of MINK1 in response to inhibition of Cullin-Ring E3 ligases (Figure 6B) our results indicate that MINK1 is regulated – similarly to β-catenin – by an APC-containing destruction complex. Importantly, increased Mink1/Msn levels after loss of WT Apc/APC1&2 were also observed *in vivo*, in mouse intestinal tissue and follicular cells in *Drosophila* (Figure 5). Consistent with its localization to cell-cell junctions, over-expression of MINK1-GFP resulted in increased cell adhesion (Figure 6D and E). This is in contrast to a previous study, in which cells overexpressing full-length MINK1 did not grow in clusters but in isolation, suggesting decreased adhesion between cells (Hu et al., 2004). However, in this case, the effects were not quantified and additional studies of MINK1 overexpression on cell-cell adhesion do not exist. It is conceivable, that enhanced cell adhesion due to elevated MINK1 expression contributes to the reduced cell migration observed in *APC* mutant tissue (Mahmoud et al., 1997; Sansom et al., 2004). Moreover, directionality of cell migration could be disturbed when MINK1 expression is deregulated in response to APC loss. Evidence for a role of mammalian MINK1 in PCP is limited (Daulat et al., 2012); however, a role for its *Drosophila* homologue Msn in epithelial PCP has been firmly established (Lewellyn et al., 2013; Horne-Badovinac et al., 2012; Paricio et al., 1999). In flies, both Apc and Msn act downstream of Dishevelled, which was described as a ‘branchpoint’ between the canonical Wnt and the non-canonical PCP pathway (Wallingford and Habas, 2005). Our results indicate that regulation of Msn by APC is conserved in flies (Figure 5C), suggesting an additional level of crosstalk between these signaling pathways. Future experiments will focus on elucidating the molecular mechanisms of MINK1 regulation by APC and the identification of downstream effectors mediating the effects of MINK1 overexpression on cell adhesion.

## Author Contributions

Conceptualization, O.P. and I.N.; Methodology, O.P., N.L., J.J. (*Drosophila* models), I.P.N., A.R.F (MINK1 KO cell lines), A.R.F (mNeonGreen-MINK1 cell lines); Formal Analysis, O.P. (lead), M. H. T. (MINK1 PTM proteomics), N.L. (live *Drosophila* imaging); Investigation, O.P. (lead), N.L. (live *Drosophila* imaging), I.P.N. (mouse experiments); Resources, M.H.T. (label-free mass spectrometry), J.A.P. (TMT mass spectrometry); Writing – Original Draft, O.P.; Writing – Review & Editing, I.N., K.M.H.; Visualization, O.P.; Funding Acquisition, O.P., K.M.H., I.N.

## Acknowledgments

We would like to acknowledge the Nikon Imaging Center at Harvard Medical School and the Dundee Tissue Imaging Facility at the School of Life Sciences, University of Dundee (supported by Wellcome grant WT101468) for providing help and equipment for microscopy imaging of cells and live *Drosophila* samples, respectively. Stocks from the Bloomington Drosophila Stock Center, which is supported by NIH grant P40OD018537, were used in this study. We would like to thank C. MacKintosh, G. Murugesan, R. Hay, E. Jaffrey, T. Kurz, and M. Keuss for sharing advice and reagents. For their help in the realization of TMT MS experiments we would like to thank S. Gygi and the Taplin Mass Spectrometry Facility at Harvard Medical School. This study was supported by a CRUK PhD fellowship to O.P.; a CRUK program grant to R. Hay supporting M.H.T.; NIH/NIDDK grant K01 DK098285 to J.A.P; a FONDECYT 11150532 and PAI79150075 grant to A.R.F.; N.L and J.J. were supported by a Sir Henry Dale fellowship (Wellcome/Royal Society: 100031Z/12/Z) to J.J.; NIH/NCI grant U01CA199252 to K.M.H.; a CRUK program grant to I.N. supporting I.P.N.

## STAR Methods

### Contact for Reagent and Resource Sharing

Further information and requests for resources and reagents should be directed to and will be fulfilled by the Lead Contact, Inke Näthke

### Experimental Model and Subject Details

#### Mice

All mice were obtained from The Jackson Laboratory and bred and maintained in accordance with their recommendations under specific pathogen-free conditions in the Biological Resource Unit at the University of Dundee. Compliant with the ARRIVE guidelines the project was approved by the University Ethical Review Committee and authorized by a project license under the UK Home Office Animals (Scientific Procedures) Act 1986.

#### Cell Lines

HeLa (female), Colo320 (female), HCT116-Haβ92 (male), U2OS (female), SW480 (male) cells were grown at 37 °C and 5% CO2 in Dulbecco’s modified eagle medium (DMEM) with 10% fetal bovine serum, 50 U/mL penicillin/streptomycin, and 1% v/v non-essential amino acids. HeLa SEC-C, U2OS SEC-C, U2OS Flp-In™ T-Rex™ and cell lines generated from these were grown as described above, with the addition of 100 μg/mL Hygromycin B and 15 μg/mL Blasticidin to the cell culture medium.

For immunoprecipitations cells were grown to 90% confluency on 15 cm tissue culture dishes.

#### Flies

Flies were reared on standard corn meal food at 25 °C.

### Method Details

### U2OS SEC-C MINK1 knockout cell lines

Analysis of the N-terminal coding region of *MINK1* (ensembl ENSG00000141503) predicted potential gRNAs with high target affinity, high efficiency and low off-target scores with binding sites in exon 1 and exon 2 using CRISPR Design. We used the best scoring gMINK1 target site in exon 1 (CGGACAGGTCGATGTCGTCC [AGG]) with a score of 95 and 12 predicted off-target sites in other genes. The gRNA sequence was cloned into pBabeD pU6 and sequence-verified. U2OS cells stably expressing Cas9 (U2OS SEC-C) were co-transfected with 3 μg pBabeD pU6 gMINK1 using Lipofectamine 2000 according to the manufacturer’s instructions. Cells were grown in DMEM supplemented with 10% FBS, 2 mM L-glutamine, and 100 μg/mL Normocin™. After 12 h, medium was replaced with fresh medium with 4 μg/mL Puromycin. After 48 h of selection, 2 μg/mL doxycycline was added to induce Cas9 expression. After 72 h, single cells were sorted with an Influx™ cell sorter (BD Biosciences) into 96-well plates containing DMEM with 20% FBS, 2 mM L-glutamine, 100 U/mL penicillin, 100 μg/mL streptomycin, and 100 μg/mL Normocin™. MINK1 protein expression was screened by western blotting. Genomic DNA of MINK1 KO cells was amplified by PCR and sequenced to confirm the introduction of frameshift mutations.

### HeLa SEC-C mNeonGreen-MINK1

The same pBabeD pU6 gMINK1 vector used for the generation of MINK1 knock-out cells was used for the fusion of mNeonGreen to the N-terminus of MINK1. A donor vector was designed to replace the ATG start codon of *MINK1* with the start codon of an mNeonGreen cDNA cassette, which was flanked by ˜500 bp homology arms to allow for homologous recombination once cleavage by CRISPR/Cas9 took place. The mNeonGreen insert was designed to remain in-frame with *MINK1* and additional silent mutations were introduced into the donor homologous right arm to prevent repeated cleavage by CRISPR/Cas9. The donor vector was synthesized by GeneArt (Life Technologies). HeLa cells stably expressing Cas9 (HeLa SEC-C) were co-transfected with 3 μg pBabeD pU6 gMINK1 and 3 μg of mNeonGreen donor vector using Lipofectamine 2000. Cells were grown for 12 h in DMEM supplemented with 10% FBS, 2 mM L-glutamine, and 100 μg/mL Normocin™. Medium was then replaced by fresh medium supplemented with 2 μg/mL doxycycline to induce Cas9 expression for 24 h. mNeonGreen expression was analyzed on a LSRFortessa™ cell analyzer or FACSCanto flow cytometer (BD Biosciences) using 488 nm excitation and 530±30 nm emission wave length. Single mNeonGreen expressing cells were sorted with an Influx™ cell sorter (BD Biosciences) into 96-well plates containing DMEM supplemented with 20% FBS, 10% Opti-MEM, 2 mM L-glutamine, 100 U/mL penicillin, 100 μg/mL streptomycin, and 100 μg/mL Normocin™. Single cell clones were expanded and expression of mNeon-MINK1 was validated by western blotting and microscopy (see also Rojas-Fernandez et al., 2015).

### U2OS Flp-In T-Rex MINK1-GFP/GFP

Stable cell lines with tetracycline/doxycyline-inducible expression of MINK1-GFP and GFP, respectively, were generated using the Flp-In™ T-Rex™ System according to the manufacturer’s instructions: U2OS Flp-In™ T-Rex™ host cells were transfected with pcDNA5 FRT/TO C-GFP or pcDNA5 FRT/TO MINK1-GFP, respectively, and pOG44, a constitutive Flp recombinase expression plasmid. Cells with successful recombination were selected by addition of Hygromycin B (100 μg/mL, resistance conferred by successful integration of the expression plasmid) and Blasticidin (15 μg/mL, resistance conferred by (already) stably integrated Tet repressor construct) to the culture medium. After three weeks, single colonies were isolated by trypsinization and expanded. Expression of MINK1-GFP and GFP, respectively, was induced by addition of 1 μg/mL tetracycline/doxycycline to the tissue culture medium.

#### Generation of fly lines and mosaic follicular epithelia

To measure Msn levels in *apc* mutants in flies, we used meiotic recombination to recombine the YFP-fused, endogenously expressed allele *msn^CPTI003908^* with the FRT[82B], *apc1*^-^, *apc2*^-^ chromosome. We then crossed *msn^CPTI003908^*, FRT[82B], *apc1*^-^, *apc2*^-^ and Hs-*flp*; Ubi-*PH^PLCδ1^::RFP*; FRT[82B], Ubi-*nls::RFP* flies and used mitotic recombination in the resulting progeny to induce *apc1-, apc2-* mutant clones in follicular epithelial cells by performing a 2 hours long, 37 °C heat-shock at a late (starting to pigment) pupal stage.

#### Cell treatment

##### siRNA transfection

Cells were seeded at a density of 2.5×10^5^ (U2OS)/ 5×10^5^ (HeLa) cells per tissue culture flask with 25 cm^2^ surface area. One day after seeding, cells were transfected with APC siRNA (equal mix of individual siRNAs #1-#3), β-catenin siRNA, or control siRNA using INTERFERin^®^ and 72 ng siRNA/flask.

##### MLN4924

HeLa cells were seeded at a density of 5×10^5^ cells per tissue culture flask with 25 cm^2^ surface area. Two days after seeding, cell culture media was replaced with media containing 3 μM MLN4924 in DMSO.

##### Doxycycline

U2OS Flp-In™ T-Rex™ cells were plated 24 h before treatment at 0.75×10^6^ cells per 10 cm tissue culture dish. To induce over-expression of GFP and MINK1-GFP media was replaced with normal growth media containing 1 μg/mL Doxycycline for 24–72 h.

#### Cell harvest

##### For protein

Cells growing in dishes/flasks were cooled on ice for five minutes, and then incubated with lysis buffer (50 mM Tris-HCl pH 7.5, 100 mM NaCl, 5 mM EDTA, 5 mM EGTA, 40 mM β-glycerophosphate, 0.5% NP-40, 1 mM sodium fluoride, 0.1 mM sodium orthovanadate, 10 μg/mL of each leupeptin, pepstatin A, and chymostatin) for five more minutes on ice. Lysates were collected by scraping, centrifuged at 14.000 rpm for 20 min at 4 °C, and supernatants were collected for further processing. For MINK1 PTMMS, cells were harvested two days after siRNA transfection with phospho-lysis buffer (270 mM sucrose, 120 mM NaCl, 50 mM NaF, 50 mM Tris-HCl pH 7.5, 10 mM β-glycerophosphate, 5 mM Na-pyrophosphate, 1% Triton X-100, 0.1% β-mercaptoethanol, 1 mM Benzamidine, 1 mM EDTA, 1 mM EGTA, 1 mM Na-orthovanadate, 1 mM PMSF, 0.1 μM Microcystin-LR, 1x cOmplete™ Protease Inhibitor Cocktail) as described above.

##### For RNA

Total RNA was isolated using the NucleoSpin^®^ RNA II Kit according to the manufacturer’s instructions.

#### Mouse intestinal tissue harvest

Mice were sacrificed by cervical dislocation and their small intestine was removed and placed into ice-cold PBS. Intestines were flushed with PBS, cut into manageable pieces and flash frozen in liquid nitrogen. A pestle and mortar were frozen overnight, placed on dry-ice and lined with extra thick aluminum foil. Frozen tissue was placed in the mortar with a small amount of liquid nitrogen and the pestle was used to grind frozen tissue into powder. Lysis buffer (see cell harvest for protein) was added to tissue powder and transferred to 1.5 mL tubes. Lysates were thawed on ice and centrifuged at 14,000 rpm for 20 min at 4°C. Supernatants were collected for further processing.

#### Immunoprecipitations

##### For AE-MS and APC Co-IPs

Per experiment, 40 μl protein G-sepharose was washed with protein lysis buffer and incubated for 12 h with 80 μg (for AE-MS)/20-40 μg (for WB) of N-APC/C-APC antibody or control antibody targeting the V5 tag at 4 °C on a rotating wheel. Antibodies were crosslinked to sepharose using 5 mM bis[sulfosuccinimidyl]suberate in 50 mM borate buffer. Crosslinking reaction was quenched with 200 mM glycine pH 2.3 for 15 minutes. Beads were washed with protein lysis buffer and, if necessary, transferred to 5 mL reaction tubes. Antibody-crosslinked sepharose was incubated with pooled cell lysates harvested from five 15 cm dishes (AE-MS and validation Co-IPs)/10 mg protein lysate (all other APC Co-IPs) for 12 hours at 4 °C on a rotating wheel. Beads were washed repeatedly with 20 mM Tris-HCl ph 7.5, 150 mM NaCl, 1 mM EDTA, 0.05% Triton X-100, and 5% glycerol. Proteins were eluted by addition of 40 μl 1.3x NuPAGE™ LDS sample buffer and incubation at 99 °C for 10 minutes with agitation.

##### For MINK1 IPs for PTM-MS analysis

Per IP one μg MINK1 antibody was bound to 25 μl protein A-sepharose for 12 h at 4 °C, and incubated with pooled cell lysates harvested from five 25 cm^2^ tissue culture flasks for 12 h at 4 °C. After washing with phospho-lysis buffer, proteins were eluted as described above for APC Co-IPs.

#### SDS-PAGE and Western Blotting

Protein samples (50 μg (cell lysates)/100 μg (tissue lysates) were separated on pre-cast NuPAGE™ 4-12% gradient Bis-Tris polyacrylamide protein gels. Proteins were transferred by wet electroblotting to nitrocellulose membranes in 48 mM Tris-HCl, 390 mM glycine, 0.02% SDS and 7.5% methanol. Transfer was performed at 4 °C at 30 V for at least 12 h. Membranes were incubated with WB blocking buffer (150 mM NaCl, 50 mM Tris-HCl pH 7.5, 5% w/v milk powder, 1% donkey serum, 0.02% Triton X-100, 0.01% sodium azide) for one hour at RT. Primary antibodies were diluted in WB blocking buffer or 5% w/v BSA, 0.1% Tween-20, 150 mM NaCl, and 50 mM Tris-HCl pH 7.5, depending on manufacturer’s instructions. Secondary antibodies were diluted 1:5000-10,000 in blocking buffer and incubated with the membrane for one hour at RT. Before and after incubation with secondary antibody, membranes were washed in 150 mM NaCl, 50 mM Tris-HCl, and 0.02% Triton X-100. Immunoblots were imaged with the Li-Cor Odyssey imaging system and signal intensity of protein bands was quantified and corrected for background signal using Image Studio Software.

#### Immunofluorescence

Cells were grown on collagen-coated No. 1.5 cover glass in 6-well tissue culture plates for several days prior to staining. Cells were fixed for 10 min with −20 °C methanol, permeabilized using 1% NP40 in PBS for 10 min, and incubated with IF blocking buffer (5% normal goat serum, 2% w/v BSA, 0.1% Triton X-100 in 1x PBS) for 30 min at RT. Cells were washed with 0.2% w/v BSA in 1x PBS, in between steps. Anti-MINK1 antibody was diluted 1:250 in 2% BSA w/v and 0.1% Triton X-100 in 1x PBS and incubated overnight at 4 °C in a humid chamber. After repeated washing, cells were incubated with 20 μg/mL Hoechst 33342 and Alexa Fluor^®^ 594 anti-rabbit antibody diluted 1:500 in 2% BSA w/v and 0.1% Triton X-100 in 1x PBS for one hour at room temperature. Cells were washed and cover slips were mounted onto microscopy slides using 90% glycerol with 0.5% N-propyl gallate.

Images were acquired with an inverted Nikon Eclipse Ti-E fluorescence microscope equipped with a Hamamatsu ORCA-R^2^ digital CCD camera and a Prior Scientific Lumen 200PRO light source, using a Plan Apo 60x NA 1.4 objective lens. Images were acquired with the MetaMorph software and without camera binning. 480/40 excitation and 535/50 emission filters were used for Hoechst and 545/30 excitation and 620/60 emission filters were used for Alexa594. Image brightness and contrast was adjusted equally for each image using Fiji software.

#### Live imaging

##### Cells

Cells were grown on 35 mm glass bottom dishes in DMEM without phenol red supplemented with 10% fetal bovine serum and 50 U/mL penicillin/streptomycin. Images were acquired with an inverted Nikon Eclipse Ti-E fluorescence microscope equipped with a Hamamatsu ORCA-R^2^ digital CCD camera and a Prior Scientific Lumen 200PRO light source, using a Plan Apo 60x NA 1.4 objective lens. Images were acquired with the MetaMorph software and with 2×2 binning. 480/40 excitation and 535/50 emission filters were used to image mNeonGreen. In addition, differential interference contrast (DIC) images were acquired. Image brightness and contrast was adjusted equally for each image, using Fiji software.

##### Drosophila egg chambers

Msn::YFP-expressing, mosaic *apc1^-^*, *apc2^-^* mutant female flies were dissected 24 hours after hatching. Ovaries were placed into a drop of glucose- (1 g/L) and insulin- (0.2 g/L) supplemented Schneider’s medium in a 35 mm glass bottom dish and dissected into individual ovarioles, by gently pulling on the germarium to dissociate them from the surrounding muscles sheath. They were then imaged on a SP8 confocal microscope (LEICA) equipped with a 63x NA 1.2 water immersion objective within the hour following dissection. The cytoplasm of *apc1^-^*, *apc2^-^* mutant cells (identified by the loss of the Nls::RFP marker) and neighboring control cells was manually segmented and used as a mask in which Msn::YFP levels were measured.

#### Cell adhesion assay

Ninety-six-well plates were coated with 10 μg/cm^2^ collagen overnight at 4 °C. Excess collagen solution was aspirated and plates were dried overnight in a sterile tissue culture hood and sterilized by UV light exposure for two hours. Collagen-coated plates were prepared just before use. Wells were washed once with PBS and then incubated for one hour with DMEM + 0.5% bovine serum albumin (BSA) at 37 °C. Meanwhile, cells were washed twice with PBS and detached by incubation with 10 mM EDTA in PBS for 10 min at 37 °C. Cells were washed twice with DMEM to remove EDTA, counted using a Cellometer^®^ Auto T4 bright field cell counter (Nexcelom Bioscience), and diluted to a density of 1×10^5^ cells/mL in DMEM + 0.1% BSA. One-hundred μl cell suspension (10,000 cells) was added to each well of coated 96-well plates (after aspiration of blocking buffer) and incubated for 1 h at 37 °C. Loosely attached cells were removed by vigorous shaking of the plate for 10 s, and washing with DMEM + 0.1% BSA. Adherent cells were fixed with 4% paraformaldehyde for 10 min at room temperature, washed, and stained for 10 min with 5 mg/mL crystal violet in 2% ethanol. Plates were washed once with water and then dried overnight. Crystal violet stain was solubilized with 200 μl 2% SDS/well for 30 min at room temperature. Fifty μl solution/well were transferred to new 96-well plates and diluted with water 1:4. Absorption was measured at 550 nm using a Synergy H1 Hybrid multi-mode microplate reader (BioTek).

#### Real Time-quantitative PCR (RT-qPCR)

Each 20 μl RT-qPCR reaction contained 0.3 μM forward and reverse primer, 10 μl PerfeCTa SYBR^®^ Green FastMix and 5 ng cDNA. cDNA was synthesized from isolated RNA using the qScript™ cDNA Synthesis Kit with one μg of total RNA per reverse transcriptase reaction and according to the manufacturer’s instructions. RT-qPCRs were performed in white 0.2 mL semi-skirted 96-well plates with a CFX Connect™ Real-Time PCR Detection System (Bio-Rad) using the following temperature program: 3 min - 95 °C; 30 s - 95 °C, 30 s – 59.3 °C, 30 s - 72 °C (40 cycles); 60 s - 95 °C, 60 s 55 °C, melt curve 55-95 °C. All reactions were performed in triplicate. CT values obtained for target genes were normalized to values obtained for *ACTB* (reference) and relative mRNA expression was calculated according to Pfaffl using the following formula:

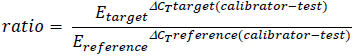

Efficiency (E) of all primers was calculated from the slope of a standard curve from a 1:10 dilution series: 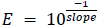 (Pfaffl, 2001). Efficiency was determined separately for each cell line: *ACTB* - HeLa E: 2.009, U2OS: 1.969; *AXIN2* - HeLa E: 2.025, U2OS: 1.962; *CTNNB1* - HeLa E: 1.943, U2OS: 1.912; *MINK1* - HeLa E: 1.995, U2OS: 2.019.

#### Protein analysis by mass spectrometry

##### Label-free MS

For Co-IPs, resin-bound proteins were eluted by boiling in 1.3x LDS sample buffer for 10 min. Co-IP and complete lysate samples were separated by SDS-PAGE on 4-12% gradient Bis-Tris polyacrylamide gels as described above and proteins were visualized by Coomassie Blue staining. To avoid mixing of high abundant immunoglobulin fragments with APC interaction partners of lower abundance in Co-IP samples, gel lanes were divided into three regions (A, B, and C). Fraction C included gel regions containing mostly heavy and light chains of co-eluted antibody. Fractions A and B contained proteins present in the upper and lower half of the gel, respectively. For MINK1 IPs, gel regions containing MINK1 were excised generously. For complete lysate samples, gel lanes were subdivided into three parts/lane. In-gel tryptic digestion was performed as described previously (Shevchenko et al., 2006). Unless stated otherwise, incubation steps were performed with shaking at room temperature and gel slices were incubated with a solution volume which covered gel slices generously. Solutions were removed from gel pieces as complete as possible using gel-loading tips, in between steps. In summary: gel slices were cut into approximately 1×1×1 mm cubes, destained in 50 mM ammonium bicarbonate (ABC) and 50% acetonitrile (ACN), and subsequently dehydrated in 100% ACN. Disulfide bonds were reduced with 10 mM dithiotreitol in 100 mM ABC for 30 min and alkylated with 50 mM iodoacetamide in 100 mM ABC for 30 min in the dark. Gel pieces were washed for 15 min in 100 mM ABC, and 15 min in 20 mM ABC and 50% ACN, before dehydration in 100% ACN. Proteins were digested ingel with trypsin in 20 mM ABC and 9% ACN at 30 °C overnight, with a protein:enzyme ratio of 20:1 (protein amounts were estimated from Coomassie Blue staining). Buffer amounts were limited to ensure complete coverage of gel slices while avoiding excess liquid. Peptides were extracted from gel pieces through successive addition of 100% ACN (30 min incubation), 5% formic acid in 50% ACN (10 min incubation, repeated once), and 100% ACN (10 min incubation). Solutions from all extraction steps were combined and dried down by vacuum centrifugation. Peptides were solubilized in 1% FA and analyzed by LC-MS/MS on a Q Exactive mass spectrometer (Thermo Scientific) coupled to an EASY-nLC 1000 liquid chromatography system via an EASY-Spray ion source (Thermo Scientific) with a 75 μm × 500 mm EASY-Spray column (Thermo Scientific) heated to 40 °C. An elution gradient duration of 240 min was used, fractionating mostly over the 3-40% acetonitrile range (further gradient details available upon request). Data were acquired in the data-dependent acquisition mode. Full scan spectra (300-1800 Th) were acquired with resolution of 70,000 at 400 Th (after accumulation to a target value of 1,000,000 with maximum injection time of 20 ms). The ten most intense ions were fragmented by higher-energy collisional dissociation (HCD) and measured with resolution 17,500 at 200 m/z and a target value of 500,000, with a maximum injection time of 60 ms. Intensity threshold was 2.1e^4^. Unassigned, +1 and >8+ charge peptides were excluded, and peptide matching was set to “preferred”. A 40 second dynamic exclusion list was applied.

##### TMT MS

Frozen cell pellets were lysed in 8 M urea and 200 mM EPPS, pH 8.5 and lysates were additionally passed ten times through a 21-gauge needle. Disulfide bonds were reduced using 5 mM tris(2-carboxyethyl)phosphine (30 min, RT) and alkylated with 10 mM iodoacetamide (30 min, RT in the dark). Alkylation reaction was quenched with 10 mM dithiotreitol for 15 min at RT. Per sample 100 μg protein (protein concentration determined prior to reduction/alkylation by BCA assay) were precipitated using methanol-chloroform precipitation and digested at RT with Lys-C protease in 200 mM EPPS, pH 8.5 at a 100:1 protein:enzyme ratio overnight. More complete protein digestion was achieved through addition of trypsin (100:1 protein:enzyme ratio) for an additional 6 h at 37 °C. Acetonitrile was added to sample to a concentration of approximately 30%, and peptides were labelled with 0.2 mg TMT isobaric label reagent per sample for one hour at RT. Labelling reactions were quenched with addition of hydroxylamine to 0.3% (v/v). Samples were combined at a 1:1:1:1:1:1:1:1 ratio and dried down by vacuum centrifugation. Excess TMT label was removed by C18 solid-phase extraction. The pooled sample was fractioned by off-line basic pH reversed-phase HPLC over a 50 min 5-35% acetonitrile gradient in 10 mM ammonium bicarbonate pH 8 into 96 fractions using an Agilent 300Extend C18 column (Wang et al., 2011). Collected fractions were combined into 24, of these 12 non-adjacent fractions were desalted using StageTips, dried by vacuum centrifugation and peptides were solubilized in 5% acetonitrile and 5% formic acid for subsequent LC-MS/MS analysis (Paulo et al., 2016). Approximately two μg of each sample were analyzed on an Orbitrap Fusion Lumos mass spectrometer (Thermo Fisher Scientific) coupled to a Proxeon EASY-nLC 1200 liquid chromatography pump (Thermo Fisher Scientific) and a 100 μm × 35 cm microcapillary column packed with Accucore C18 resin (2.6 μm, 150 Ã, Thermo Fisher). Peptides were fractionated over a 150 min gradient of 3 – 25% acetronitrile in 0.125% formic acid. An MS^3^-based TMT method was used, as described previously (Ting et al., 2011; McAlister et al., 2014; Paulo et al., 2016). MS^1^ spectra were acquired with a resolution of 120,000, 350-1400 Th, an automatic gain control (AGC) target of 5e^5^, and a maximum injection time of 100 ms in the Orbitrap mass analyzer. The ten most intense ions were fragmented by collision-induced dissociation (CID) and analyzed in a quadrupole ion trap with AGC 2e^4^, normalized collision energy (NCE) 35, q-value 0.25, maximum injection time 120 ms, and an isolation window of 0.7 Th. MS^3^ spectra were acquired in the Orbitrap mass analyzer (AGC 2.5e^5^, NCE 65, maximum injection time 150 ms, 50,000 resolution at 400 Th) after fragmentation of MS2 ions by HCD. Isolation windows were chosen depended on charge state z (z=2 1.3 Th, z=3 1 Th, z=4 0.8 Th, z=5 0.7 Th).

#### Raw MS data analysis

Raw MS data files were processed using MaxQuant with the integrated peptide search engine Andromeda (Cox et al., 2011) using default settings. MS/MS spectra were searched against the UniProt human proteome sequence database. Accurate label-free quantification and normalization were achieved by implementation of the MaxLFQ algorithm within the MaxQuant software environment, applying a minimum ratio count of 2 (Cox et al., 2014). For label-free samples the ‘match between runs’ option with default settings was enabled. TMT-labelled samples were quantified by reporter ion MS^3^ – TMT10plex (Lys & N-terminal 126C-130N), with a reporter mass tolerance of 0.003 Da.

Along with standard modification settings Acetyl (K), GlyGly (K), and Phospho (STY) were searched as variable post-translational modifications on MINK1. One percent FDR filtering was applied at protein, peptide, and site levels.

#### MS data processing

Further MS data analysis was performed using Perseus. Proteins identified by one post-translationally modified peptide only (‘only identified by site’), proteins identified by peptide from the decoy database (‘reverse’), and all non-human contaminants and human contaminants, except cytoskeletal components, were filtered out. Data were then log2 transformed.

The filtered APC AE-MS data set contained 5,571 identified proteins, of which 5,521 were measured, i.e. a LFQ intensity value was reported by MaxQuant. From these only proteins measured in all four replicates of at least on IP with N-APC, C-APC or control antibody, respectively, were carried forward (4,016 proteins). Missing values were imputed from a normal distribution using standard settings (width: 0.3 × standard deviation of measured values, down shift: 1.8 in units of standard deviation of measured values).

The filtered label-free proteome data set contained 5,982 identified proteins, of which 4,927 were measured in at least three replicates of at least one condition and only these were used for further analysis. Missing values were imputed from a normal distribution using standard settings (see above).

The filtered TMT proteome data set contained 6,949 identified proteins, of which 6,923 were measured in all analyzed samples. Only these proteins were used for further analysis.

Statistical tests were performed as indicated in the figure legends using standard settings unless stated otherwise.

Enrichment analysis of category terms within the group of potential APC interactors identified by AE-MS (171 proteins) relative to all proteins measured in this experiment (4,016 proteins) was calculated by Fisher Exact Test using default settings with a Benjamini-Hochberg FDR <0.02 used for truncation.

For MINK1 PTM analysis, raw peptide intensities were normalized to MINK1 protein intensity in each sample. Only peptides with PTMs that provided intensity values in all replicates of a single condition were carried forward for analysis. Zero intensity values were replaced in Perseus using default settings.

#### Network generation

The APC interaction network was generated in Cytoscape by importing interaction partners of novel (identified in this study) and known (listed in IntAct Molecular Interaction Database and/or BioGRID interaction repository) APC-binding proteins (Orchard et al., 2014; Stark, et al., 2006). Corresponding links between them were also included using information from IntAct. Interactions within IntAct are scored based on how often an interaction has been reported in the literature and which detection methods were used. Scores range from zero to one, a score of one reflecting the highest confidence interaction in the database. To disentangle the generated ‘hairball’ network, low-confidence links (IntAct MI score <0.6) were deleted. Individual nodes detached from the network and indirect APC interactors with less than two connections were omitted. The network layout was generated using the Weak Clustering algorithm and the IntAct MI score for edge weighting within the Cytoscape Allegro Layout App.

## Quantification and Statistical Analysis

Names of applied statistical tests including definitions of significance are indicated for individual experiments in the figure legends.

## Supplemental Information

**Figure S1.**
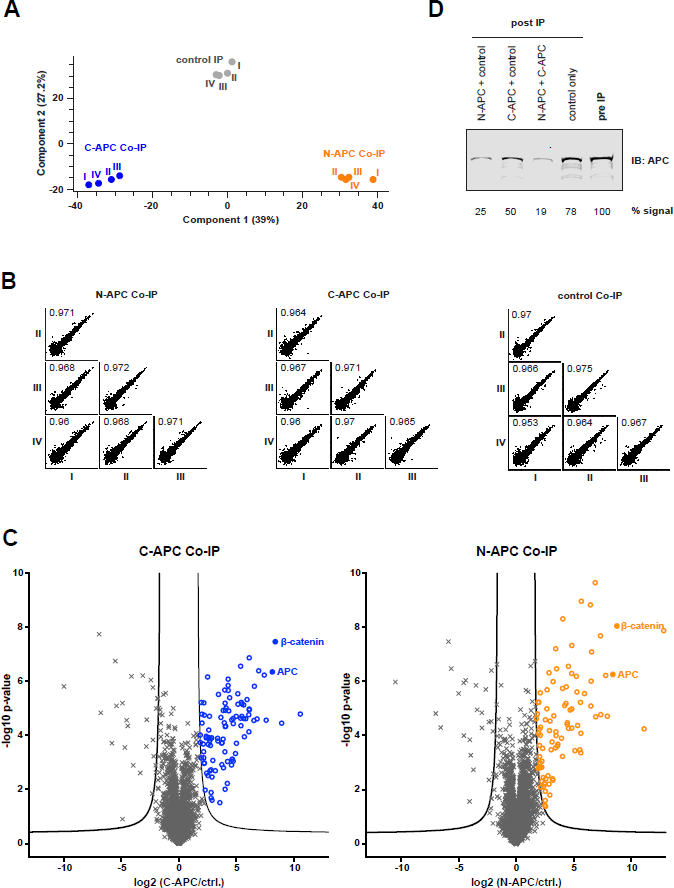
APC Affinity enrichment-mass spectrometry (AE-MS). See also Figure 1B. **A** Projections – Principal component analysis of N-APC, C-APC and control Co-IP samples. **B** LFQ intensities for all measured proteins in respective replicates plotted against each other. The Pearson correlation coefficient is indicated for each comparison. **C** Volcano plots of proteins measured in C- and N-APC Co-IPs, respectively. Plotted is the log2 fold change in mean LFQ intensities between specific APC IP and control IP (*x*-axis) against the – log p value obtained by Student’s t-test (*y-*axis). Significant enrichment in specific APC IP vs. control IP (shown as colored circles, C-APC Co-IP= 101, N-APC Co-IP= 91) was determined by two-sided t-test with permutation-based FDR <0.01 and s0 = 2 used for truncation (black line). **D** Levels of APC protein present in HeLa cell lysates after immunoprecipitation (post-IP) compared to protein levels before IP (pre-IP) using ˜28 mg protein/IP (this is equivalent to the amounts used for the APC AE-MS experiment). The pool of APC immunoprecipitated by C-APC AB (50% of total APC) overlapped with but was not equivalent to the APC pool immunoprecipitated by N-APC AB (75% of total APC), as both ABs together immunoprecipitated over 80% of total APC protein.

**Figure S2.**
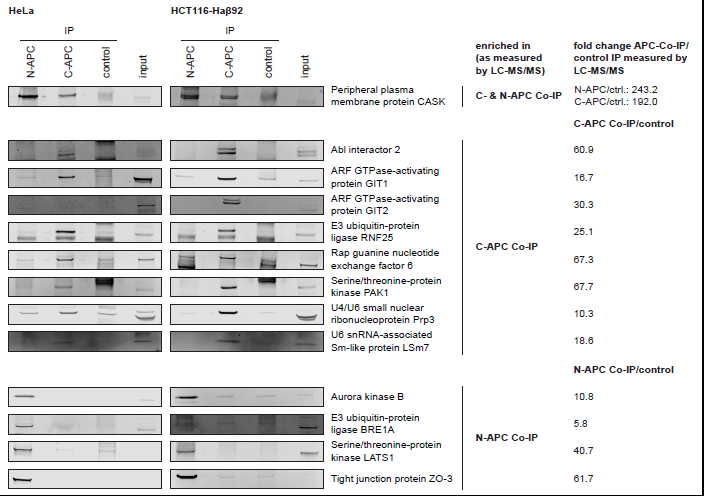
Validation of APC interactors identified by AE-MS in HeLa and HCT116-Haβ92 cells by Co-IP and WB. Input lanes: 50 μg protein. See also Figure 1B.

**Figure S3.**
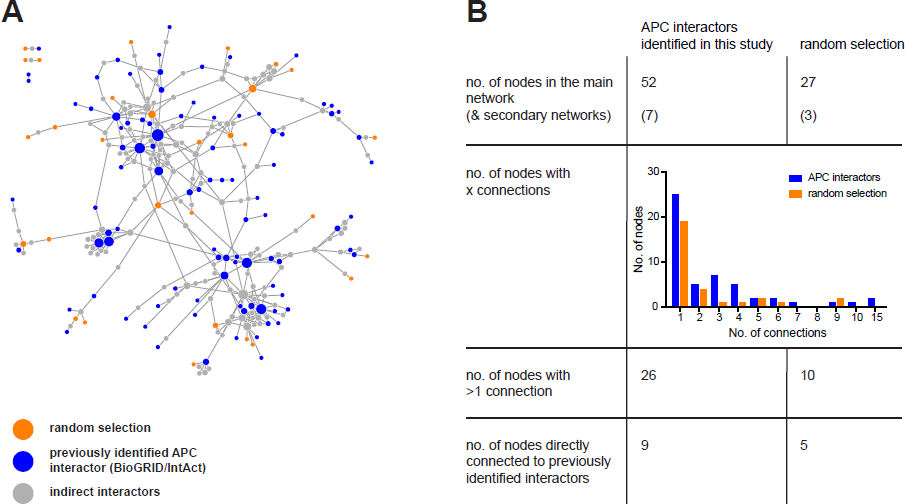
Control APC interactome network. See also Figure 2. **A** Network integrating known APC binding partners (blue), random selection (orange), and indirect interactors (grey). APC interaction partners. **B** Degree of integration into a network of known APC interactors between APC-binding proteins identified here, and the random selection in A.

**Figure S4.**
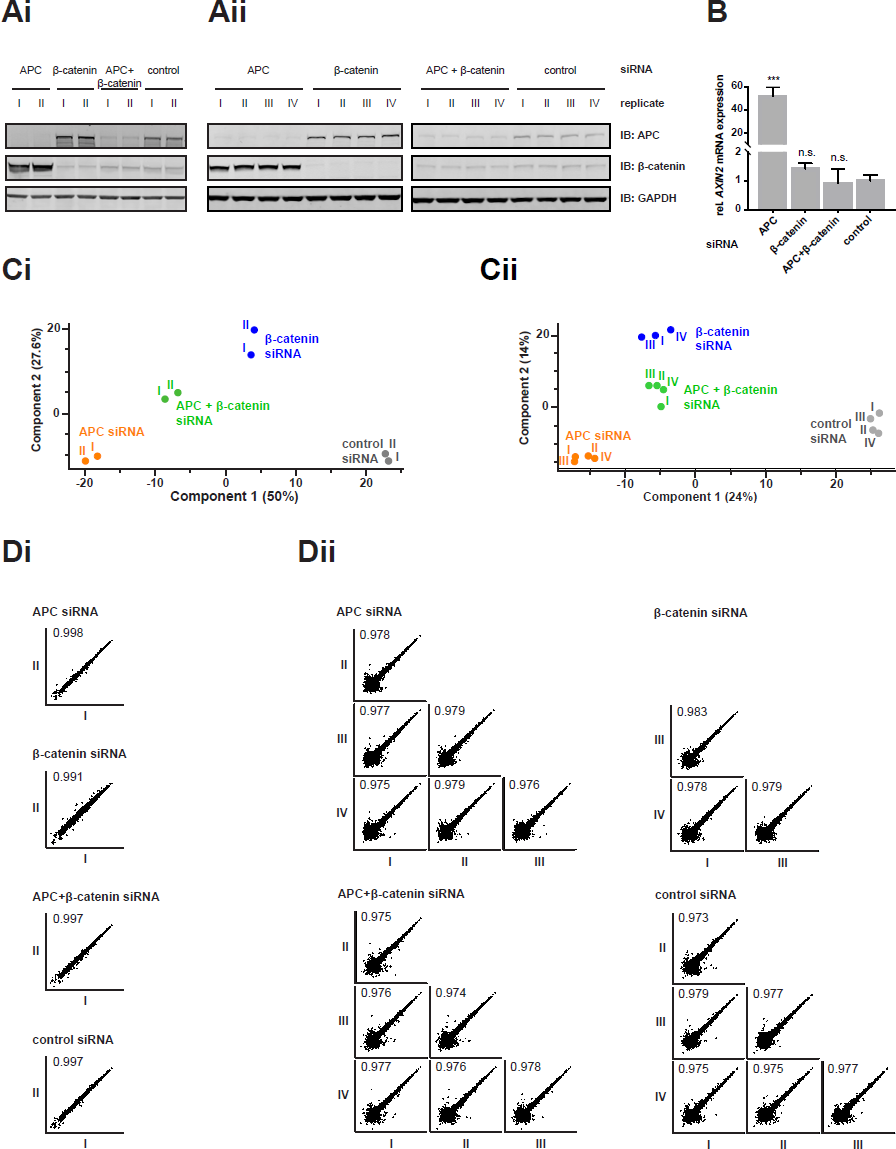
Changes in protein abundances in response to APC depletion. See also Figure 3. **A** Confirmation of efficient siRNA knockdown in MS samples by WB. Transfections were performed in duplicate for TMT (**Ai**) and in quadruplicate for label-free (**Aii**). **B** mRNA expression of the β-catenin target gene *AXIN2* analyzed by RT-qPCR. Indicated are mean expression levels relative to *ACTB* expression with SD from four independent transfections. Significance determined by one-way ANOVA followed by Dunnett’s multiple comparison test; p value: *** < 0.001. Note, this data is also shown in Figure 6A. **C** Projections - PCA of samples analyzed by TMT (**Ci**) and label-free (**Cii**) MS. **Di**: TMT MS, **Dii**: label-free MS. Sum TMT intensities/LFQ intensities for all measured proteins in respective replicates plotted against each other. The Pearson correlation coefficient is indicated for each comparison. Replicate II of the label-free β-catenin siRNA-treated sample was lost during sample processing.

**Figure S5.**
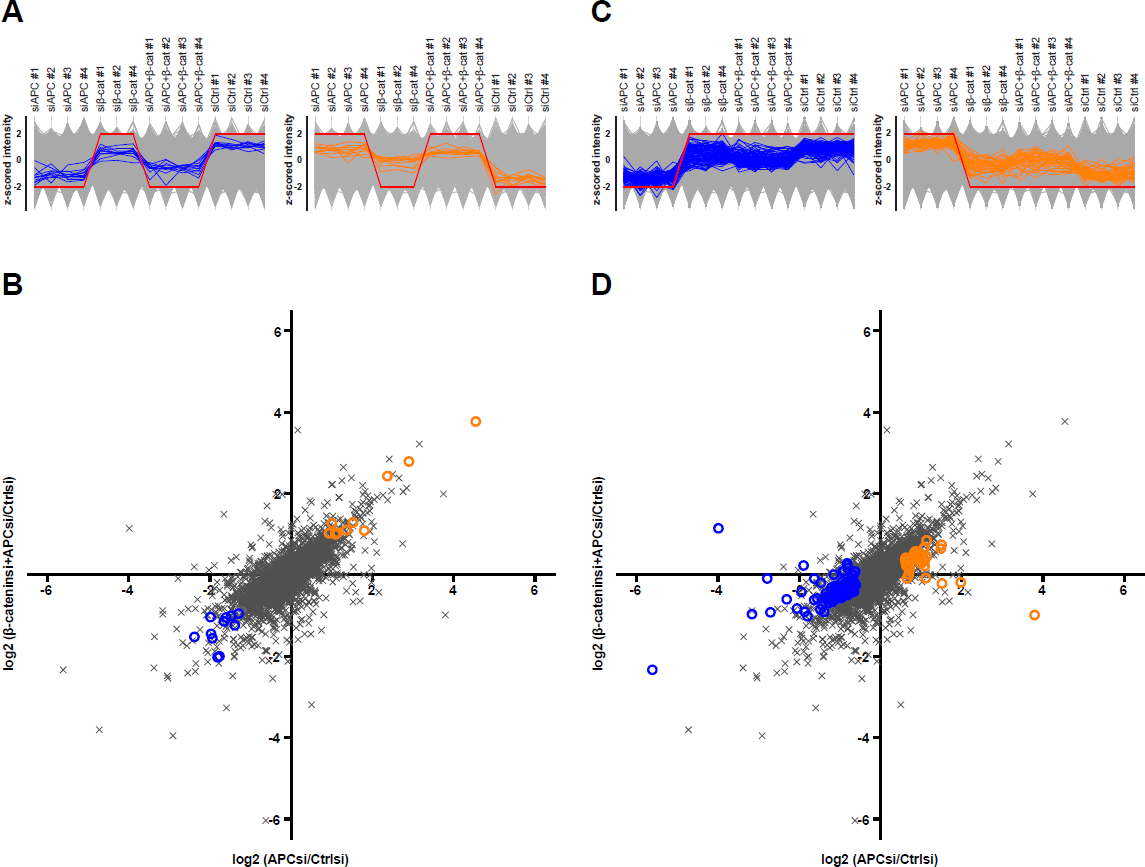
β-catenin-dependent and – independent APC targets identified by label-free MS. See also Figure 3. **A** Profiles of z-scored LFQ intensities of all measured proteins across samples. Protein identified as negative and positive β-catenin-independent APC targets are shown in orange and blue, respectively. Red lines show profiles for hypothetical ‘ideal’ targets that increase/decrease in response to APC depletion, but irrespective of change in βcatenin levels. **B** Log2 fold change in mean LFQ intensities between APC siRNA and control siRNA treated samples (*x*-axis) plotted against the log2 fold change in mean LFQ intensities between β-catenin+APC siRNA and control siRNA treated samples (*y-*axis). Proteins selected based on their LFQ intensity profiles in **A** are shown in orange and blue, respectively. **C** and **D** Same as A, but showing β-catenin-dependent APC targets.

**Figure S6.**
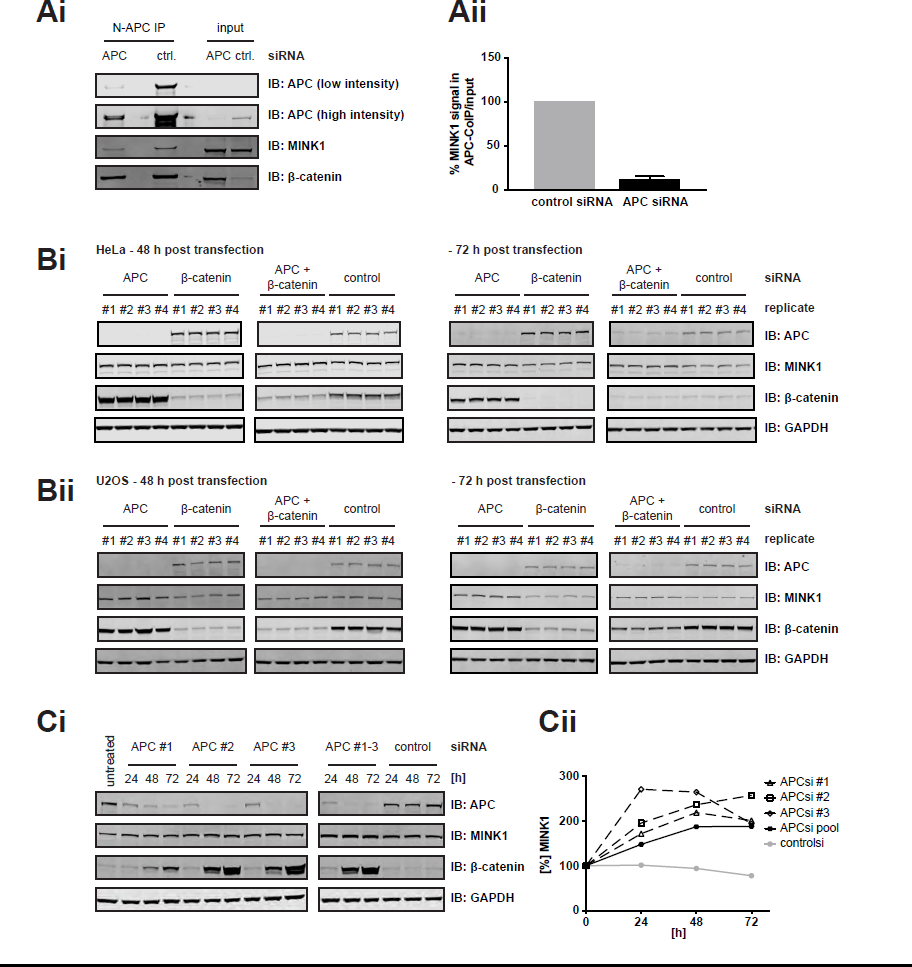
MINK1 binds to and is negatively regulated by APC. See also Figure 4. **Ai** APC Co-IPs in HeLa cells transfected with APC and control siRNA, respectively, confirming a specific interaction between MINK1 and APC. **Aii** Quantification of MINK1 WB signal in N-APC IP vs. input samples from control and APC siRNA treated cells. Shown are the mean and SD from two independent experiments. **B** Measurements of MINK1 proteins levels in response to siRNA-mediated depletion of APC and/or βcatenin in HeLa (**i**) and U2OS (**ii**) cells. Note, HeLa 72 h post transfection samples were also used for label-free total proteome measurements (see Figure S4A ii for partial blot shown here). **Ci** Measurement of MINK1 protein levels in HeLa cells transfected with individual APC siRNAs (APC siRNA #1, #2, #3), a combination of these, or control siRNA. **Cii** Quantification of MINK1 WB signal in Ci, levels are indicated relative to untransfected sample.

**Figure S7.**
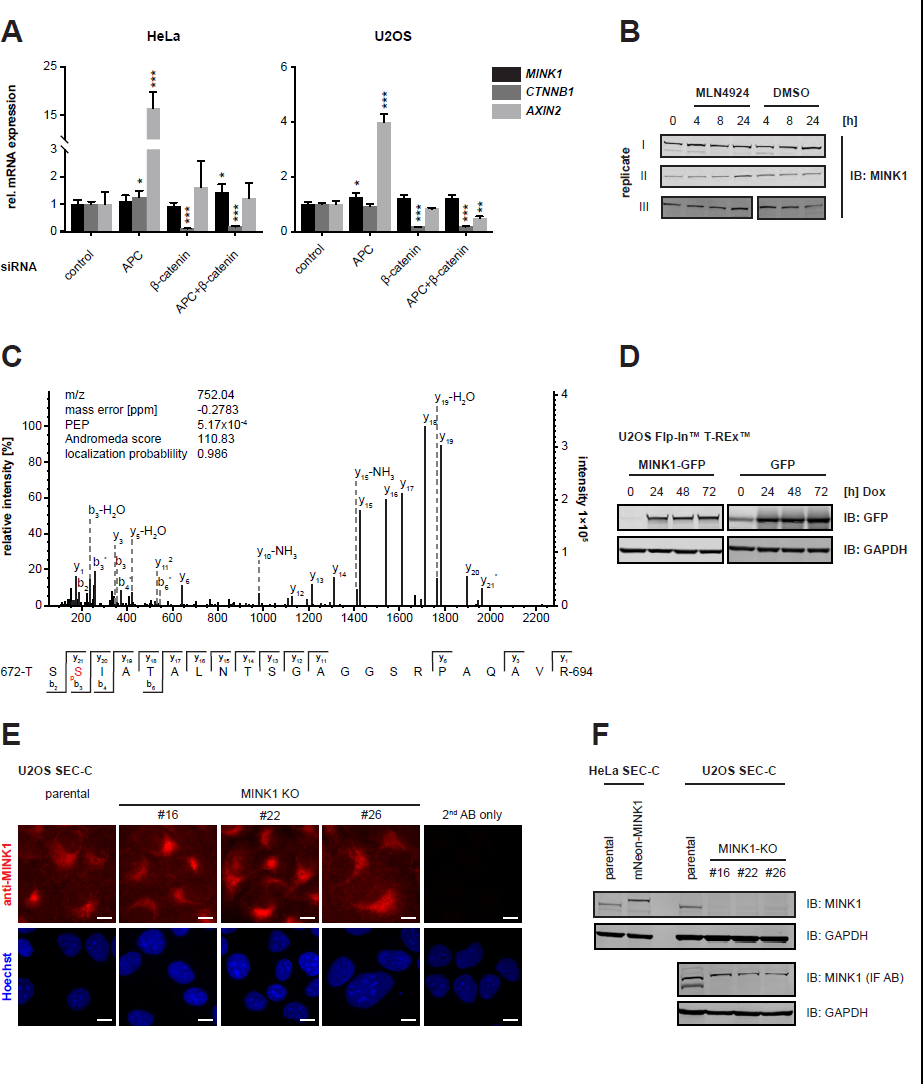
Transcriptional and post-translational regulation of MINK1 by APC and MINK1 subcellular localization. See also Figure 6. **A** mRNA expression of *MINK1*, *CTNNB1*, and *AXIN2* measured by RT-qPCR 48 h after siRNA transfection. Indicated are mean expression levels relative to *ACTB* expression with SD from four independent transfections. Significance determined by one-way ANOVA followed by Dunnett’s multiple comparison test; p value: * < 0.05, ** < 0.01, *** < 0.001. **B** MINK1 protein levels in HeLa cells after treatment with neddylation inhibitor MLN4924 [3 μM]. **C** Annotated MS/MS spectrum of the tryptic MINK1 peptide indicating phosphorylation at S647. Asterisks indicate loss of H3PO4 from the fragment ion. **D** Doxycycline-inducible overexpression of MINK1-GFP and GFP, respectively, in stable U2OS Flp-In™T-REx™ cell lines. **E** Anti-MINK1 immunofluorescence signal in fixed U2OS SEC-C parental and MINK1 knock-out cells. Scale bars: 10 μm. **F** Confirmation of tagged MINK1 expression in mNeonGreen-MINK1 HeLa SEC-C cells (left) and successful knock-out of MINK1 in U2OS SEC-C MINK1-KO clones #16, #22, and #26 (right). The MINK1 antibody used for IF in E (MINK1 (IF AB)), recognizes protein bands of similar molecular weights to MINK1 in U2OS SEC-C parental and MINK1-KO cells.

Table S1. Proteins measured by APC affinity enrichment-mass spectrometry. Related to Figure 1, Figure 2, Figure S1, and Figure S3.

Table S2. APC interactors – Enrichment Analysis. Related to Figure 1.

Table S3. β-catenin-independent and – dependent APC targets identified by TMT label mass spectrometry. Related to Figure 3, and Figure S4.

Table S4. β-catenin-independent and – dependent APC targets identified by label-free mass spectrometry. Related to Figure 3, and Figure S4.

Table S5. MINK1 post-translational modifications identified by label-free MS in APC and control siRNA-treated HeLa cells. Related to Figure 6C.

Network S1. Integrated APC interaction network. Related to Figure 2 and S3.

